# Heme oxygenase-1 deficiency affects bone marrow niche and triggers premature exhaustion of hematopoietic stem cells

**DOI:** 10.1101/492850

**Authors:** Krzysztof Szade, Monika Zukowska, Agata Szade, Witold Nowak, Maciej Ciesla, Karolina Bukowska-Strakova, Gunsagar Singh Gulati, Neli Kachamakova-Trojanowska, Anna Kusienicka, Elisa Einwallner, Jacek Kijowski, Szymon Czauderna, Harald Esterbauer, Irving Weissman, Jozef Dulak, Alicja Jozkowicz

## Abstract

While intrinsic changes in aging hematopoietic stem cells (HSCs) are well-characterized, it remains unclear how hematopoietic niche affects HSC aging. Here, we demonstrate that cells in the niche — endothelial cells (ECs) and CXCL12-abundant reticular cells (CARs) — highly express the heme-degrading enzyme, heme oxygenase 1 (HO-1), but then decrease its expression with age. RNA-sequencing shows that ECs and CARs from HO-1-deficient animals (HO-1^-/-^) produce less hematopoietic factors. Consequently, HSCs from young HO-1^-/-^ animals lose quiescence and regenerative potential. Young HO-1^-/-^ HSCs exhibit features of premature aging on the transcriptional and functional level. HO-1^+/+^ HSCs transplanted into HO-1^-/-^ recipients exhaust their regenerative potential early and do not reconstitute secondary recipients. In turn, transplantation of HO-1^-/-^ HSCs to the HO-1^+/+^ recipients recovers the regenerative potential of HO-1^-/-^ HSCs and reverses their transcriptional alterations. Thus, HSC-extrinsic activity of HO-1 prevents HSCs from premature aging and may restore the function of aged HSCs.

## Introduction

Hematopoietic stem cells (HSCs) maintain the production of all blood cells throughout life, but their functional status changes with age [1–4]. As shown both in mouse and human, aged HSCs expand their pool, however, their regenerative potential is impaired [2–5]. HSCs isolated from old mice and old humans reconstitute hematopoiesis worse than young HSCs upon transplantation, preferentially differentiate toward myeloid lineage, and accumulate DNA damage [3–7]. Age-related changes in HSCs result in reduced immunity in elderly individuals and are linked to cardiovascular diseases, myelodysplasia, and leukemic malignancies [8–11]. While the effect of aging on HSCs is now better characterized, the questions of how these changes arise and whether or not they are reversible remain yet largely unanswered [12–14].

Until now, the mechanisms found to contribute to the aging of HSCs have been mostly intrinsic to the HSCs [6,15]. These include age-related accumulation of mutations and cell-autonomous changes in the transcriptome and epigenome of HSCs [6,7,16–19]. Although HSC-extrinsic factors from the local bone marrow (BM) environment of HSCs – the HSC niche – are critical for HSCs maintenance [20–22], little is known about their contribution to HSC aging.

Recent findings indicate that HSCs occupy a perivascular niche and localize in direct proximity to endothelial cells (ECs) and mesenchymal stromal cells (MSCs) surrounding vessels [23,24]. Among the many cell types in the HSC niche, the ECs and MSCs constitute the main source of stromal cell-derived factor 1a (SDF-1a) and stem cell factor (SCF) – extrinsic factors critical for HSC maintenance [25–28]. Specific deletion of either *Sdf1 α* or *Scf* in either ECs or MSCs causes hematopoietic collapse or triggers over-activation of HSCs and their release from the niche [26,25,27,22].

Given the crucial role of the perivascular niche in maintaining HSCs, we hypothesized that HSC-extrinsic factors that support function of endothelial cells and regulate the activity of hematopoietic mediators may be implicated in HSC aging. This led us to heme oxygenase 1 (HO-1), a free heme-degrading enzyme, as a potential niche-dependent factor that may affect HSC homeostasis.

HO-1 is an antioxidative, antiinflammatory and antiapoptotic protein, undetectable in most cell types in a steady state but induced under the stress conditions [29]. Only in some cell types, as Kupffer cells in the liver or CD4+CD25+ regulatory T cells, HO-1 is constitutively expressed [30]. HO-1 deficiency disturbs iron metabolism and redistribution leading to microcytic anemia [31]. We and others showed that beyond its classical role in acute stress responses, HO-1 is important for SDF-1a signaling [32] and proper function of endothelial cells [33,34]. Here, we identified cell populations constitutively expressing HO-1 in the bone marrow niche. Using transplantation and genetic models combined with transcriptional profiling, we demonstrated that HO-1 in the niche prevents HSCs from premature aging.

## Results

### Bone marrow endothelial and stromal cells express heme oxygenase-1 in steady-state conditions

We first determined the distribution of HO-1 in the murine BM niche under steady-state conditions. Confocal microscopy analysis of mouse tibias and femurs revealed a high level of HO-1 protein in endomucin-positive (endomucin+) capillaries in the bone metaphysis (Figure 1A, Figure S1A), while HO-1 expression in endomucin+ sinusoids in the bone diaphysis, although detectable, was lower (Figure S1B). Further characterization showed that HO-1 was expressed in both endomucin+CD31+ small capillaries (Figure 1B) and bigger endomucin^-/low^CD31+ arteries (Figure 1C).

**Figure 1.**
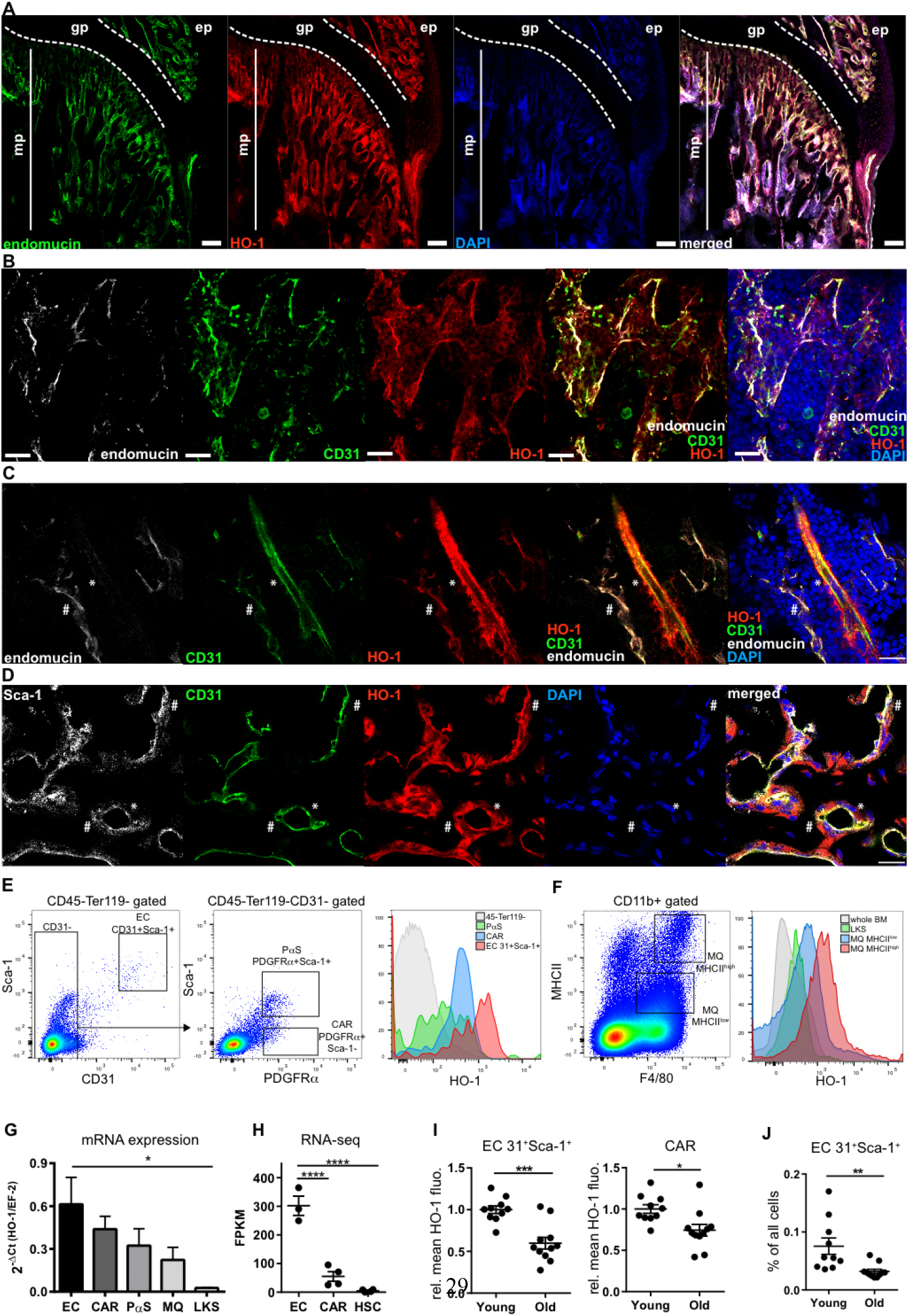
HO-1 is expressed in BM endothelial cells and pericytes. (A) Metaphysis region in the BM is reach in endomucin+ capillaries expressing HO-1. mp - metaphysis; gp - growth plate; bar 100 μm. (B) The HO-1 positive small capillaries in metaphysis express endomucin and CD31. Shown maximum intensity projection, bar 20 μm. (C) HO-1 is expressed by smaller endomucin+CD31+ capillaries (#), as well as, in bigger endomucin^-/low^CD31+ arteries (*). CD31^-^ pericytes wrapping the artery also express HO-1 (*); bar 20 μm. (D) HO-1-positive capillaries in the metaphysis expressed CD31 and Sca-1. The capillaries are enveloped by HO-1 expressing pericytes. Part of the HO-1+ pericytes express Sca-1 (#) while others show no or low Sca-1 signal (*); bar 20 μm. (E) Flow cytometry analysis revealed the highest expression of HO-1 in CD31^+^Sca-1^+^ ECs. CAR and PaS populations also express HO-1, while most of non-hematopoietic CD45^-^Ter119^-^ are HO-1-negative in steady-state conditions. (F) BM macrophages (MQ) express HO-1. The MHCII^high^ MQ express higher levels of HO-1 than MHCII^low^ MQ. Cells within whole HSPC compartment (LKS) express no or low levels of HO-1 in comparison to MQ. (G) HO-1 expression on mRNA level quantified by qPCR or (H) RNAseq levels. qPCR analysis based on two independent experiments n = 10-11/group, two-tailed t-test for two groups comparison, and one way ANOVA with Bonferroni post-test for multiple groups comparison. (I) ECs and CAR from old animals (11-12 months) express lower levels of HO-1 protein. Two independent experiments, n = 5-10 /group. (J) Old animals have lower frequency of ECs. Two independent experiments, n = 10-11/group.

We noticed that pericytes surrounding the capillaries also expressed HO-1 (Figure 1C,D). Most HO-1-expressing pericytes were PDGFRβ-positive and were present in both the metaphyseal and diaphyseal region of the bones (Figure S1C). The HO-1-expressing pericytes were also heterogeneous by the Sca-1 expression, ranging from Sca-1 positive to Sca-1 low and negative (Figure 1C, Figure S1D). Moreover, throughout the BM cavity, we detected PDGFRβ-positive cells that expressed HO-1 and produced SDF-1a (Figure S1E).

Next, we quantified the expression of HO-1 in the various bone marrow populations by flow cytometry and real-time PCR. Consistent with immunochemical staining, we identified a CD45^-^ Ter119^-^CD31^+^Sca-1^+^population of endothelial cells and two subsets of CD45^-^Ter119^-^CD31^-^ PDGFRα^+^ cells that differ in Sca-1 expression by flow cytometry (Figure 1E). Bone marrow cells with the phenotype CD45^-^Ter119^-^CD31^-^PDGFRα^+^Sca-1^-^ correspond to the previously described Cxcl12-abundant reticular cells (CARs) [28,35], while cells with the CD45^-^Ter119^-^CD31^-^ PDGFRα^+^Sca-1^+^ phenotype correspond to the PDGFRα- and Sca-1-expressing mesenchymal cells (PαSs) [36]. The flow cytometric analysis revealed that ECs, CARs, and PαSs cells all highly express HO-1 in a steady state, with the highest expression found in the CD31^+^Sca-1^+^ ECs (Figure 1E). In contrast, the vast majority of CD45^-^Ter119^-^ non-hematopoietic bone marrow cells did not express HO-1 (Figure 1E). We also examined HO-1 in BM hematopoietic cells. Among them, the highest HO-1 expression was found in macrophages (MQ), particularly in the CD11b+F4/80+MHCII^high^ fraction, followed by the CD11b+F4/80+MHCII^low^ fraction (Figure 1F). The pool of hematopoietic stem and progenitor cells (HSPC), defined as Lin^-^c-Kit^+^Sca-1^+^(LKS), expressed only low levels of HO-1 (Figure 1F).

To directly compare the expression of HO-1 between the various populations, we sorted CD31^+^Sca-1^+^ ECs, CARs, PαSs, BM macrophages (CD11b+F4/80+MHCII+), and HSPCs (Lin^-^ Kit^+^Sca-1^+^, LKS) and measured the mRNA levels of HO-1 by qPCR (Figure 1G). Concordantly with results obtained by flow cytometry, HO-1 mRNA expression was the highest in ECs, high in CARs, PαSs, and MQ, and lowest in HSPCs (Figure 1G). We also checked the expression of HO-1 in our RNA-sequencing (RNA-seq) data sets that include strictly defined sorted LT-HSCs (LKS CD150+CD48^-^CD34^-^) and the two most highly HO-1-expressing populations in the BM, namely ECs and CARs. We found that LT-HSCs expressed negligible levels of HO-1 compared to ECs and CARs (Figure 1H).

Next, we asked whether the expression of HO-1 in ECs and CARs changes with age. Using flow cytometry, we observed that HO-1 protein level was significantly lower in BM CD31^+^Sca-1^+^ ECs and CARs isolated from old animals (11-12 months old) as compared to young animals (1.5-3 months old) (Figure 1I). Moreover, the frequency of CD31^+^Sca-1^+^ECs in BM was also decreased in old animals (Figure 1J). Altogether, we found that HO-1 is highly expressed under steady-state conditions in cells that constitute the perivascular BM niche (ECs and CARs), and that its expression in the niche decreases with age.

### Lack of HO-1 dysregulates expression of hematopoietic factors by ECs and CARs in BM

Given the high HO-1 expression in ECs and CARs, we investigated how lack of HO-1 affects these populations. To this aim, we sorted ECs and CARs from wildtype (HO-1^+/+^) and HO-1 deficient mice (HO-1^-/-^), and compared their transcriptional profile by RNA-seq. DESeq2 analysis identified 111 differentially expressed genes (DEGs, FDR < 0.1, Figure 2A, Table S1 and S2), that distinguished the HO-1^-/-^ and HO-1^+/+^ ECs by hierarchical clustering and principal component analysis (PCA) (Figure 2B,C). Gene set enrichment analysis (GSEA) indicated that lack of HO-1 affects not only endothelial biology (angiogenesis, endothelial cell migration, and proliferation), but also dysregulates genes related to hematopoiesis (Figure 1D, E). Among the downregulated genes were *Scf* (*Kitl*, padj = 0.0007) and *Sdf1* (*Cxcl12*, padj = 0.06). The other dysregulated genes were implicated in integrin-mediated adhesion and signaling pathways (Figure 2D,F). Among HO-1^-/-^ CARs, we found 100 significant DEGs (Figure 2G, Table S2), that also separate HO-1^-/-^ and HO-1^+/+^ CARs by hierarchical clustering and PCA analysis (Figure 2H,I). GSEA indicated that these dysregulated genes are related to mesenchymal differentiation (osteoblast proliferation, bone development, ossification) (Figure 2J,K), hematopoietic progenitor differentiation, hematopoietic stem cell proliferation, and myeloid cell differentiation (Figure 2J,L). The remaining significantly enriched gene sets for HO-1^-/-^ vs. HO-1^+/+^ CARs also included cell adhesion and integrin signaling pathway, as well as genes involved in the regulation of endothelial migration, proliferation, and angiogenesis (Figure 2J,L).

**Figure 2.**
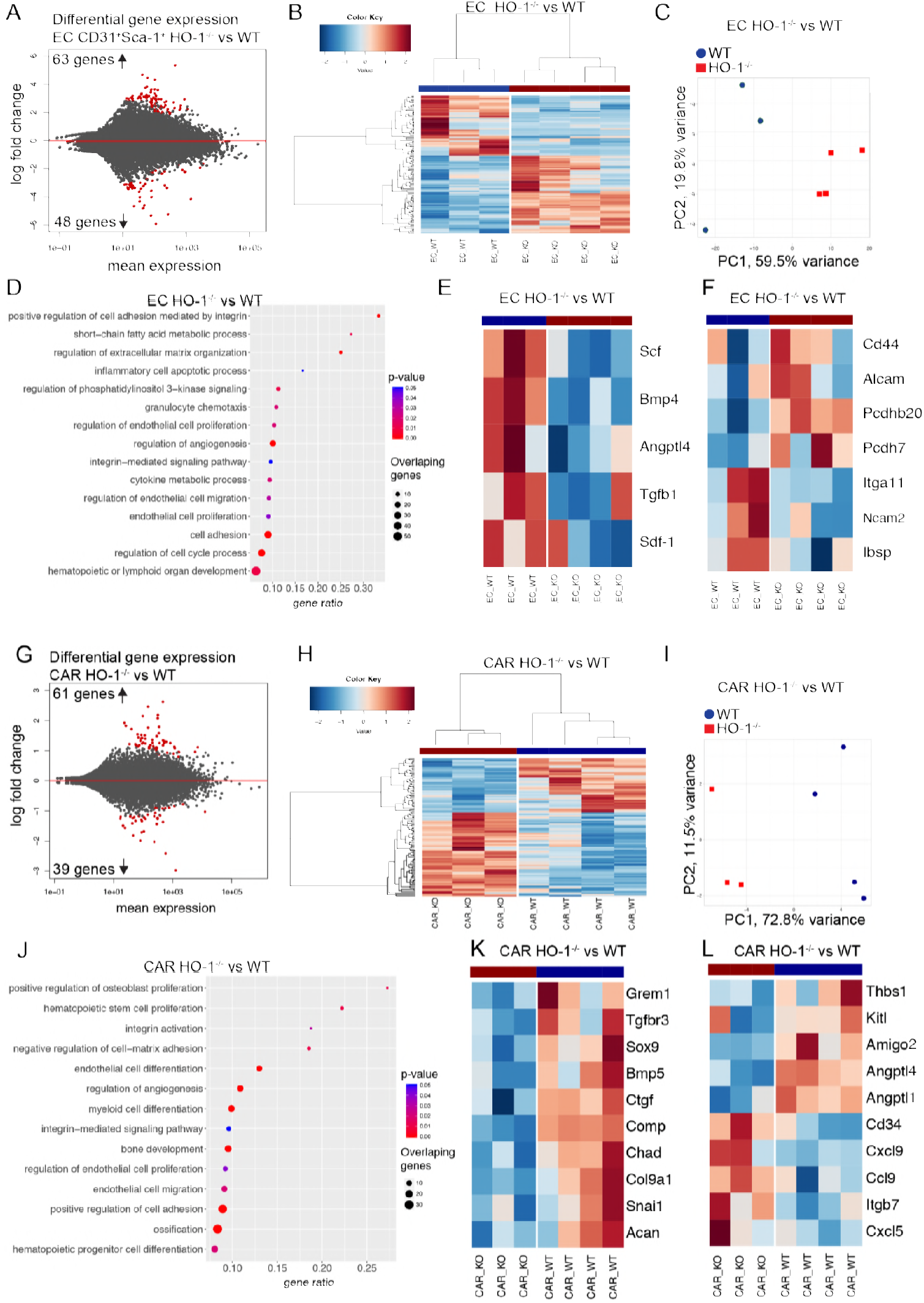
HO-1 deficiency affects expression of hematopoietic factors in ECs and CARs. (A) RNA-seq revealed 111 DEGs in HO-1^-/-^ ECs. (B) The identified DEGs separated HO-1^-/-^ and HO-1+/+ ECs by hierarchical clustering and (C) PCA. (D) Selected BP terms enriched in GSEA analysis in HO-1^-/-^ ECs. (E) Heatmap of selected hematopoietic factors and (F) adhesion molecules expression in HO-1^-/-^ and HO-1+/+ ECs. (G) RNA-seq revealed 100 DEGs in HO-1^-/-^ CARs. (H) The identified DEGs separated HO-1^-/-^ and HO-1+/+ CARs by hierarchical clustering and (I) PCA. (J) Selected BP terms enriched in GSEA analysis in HO-1^-/-^ CARs. (K) Heatmap of selected hematopoietic factors and (L) adhesion molecules expression in HO-1^-/-^ and HO-1+/+ CARs. Correlation similarity metric and average linkage clustering were used in the presented heatmaps.

### Young HO-1^-/-^ mice possess expanded pool of activated HSC

HO-1 deficiency in ECs and CARs alters the expression of niche-derived factors necessary for HSC function (Figure 2). Therefore, we investigated how HO-1 deficiency affects LT-HSCs and downstream progenitors (Figure 3A). Young (1.5-3 months old) HO-1^-/-^ mice possess increased frequency and total number of LT-HSCs (LKS CD150^+^CD48^-^CD34^-^, Figure 3B), ST-HSCs (LKS CD 150^mid^CD48^-^CD34+, Figure 3C) and MPPs (LKS CD150^-^CD48+, Figure 3D) compared to young HO-1^+/+^ mice. However, in old mice (12 months old), the frequency and total number of LT-HSCs, ST-HSCs, MPPs did not differ between HO-1^-/-^ and HO-1^+/+^ genotypes (Figure S2). Next, we determined the cell cycle status of the LT-HSCs isolated from HO-1^-/-^ mice. The flow cytometry analysis based on Ki67 and DNA content (Figure 3E) revealed that there were significantly more LT-HSCs in G1 and S/G2/M phases in young HO-1^-/-^ mice than in young HO-1^+/+^ mice (Figure 3F). Notably, this effect was specific to LT-HSCs, as young HO-1^-/-^ MPPs showed the same cell cycle status as young HO-1^+/+^ MPPs (Figure 3G). Among older mice, we still observed a higher percentage of LT-HSCs in G1 phase, although there was no difference in the frequency of LT-HSCs actively cycling in S/G2/M phases (Figure 3H). We also did not observe any differences in cell cycle status among MPPs in older HO-1^-/-^ animals (Figure 3I).

**Figure 3.**
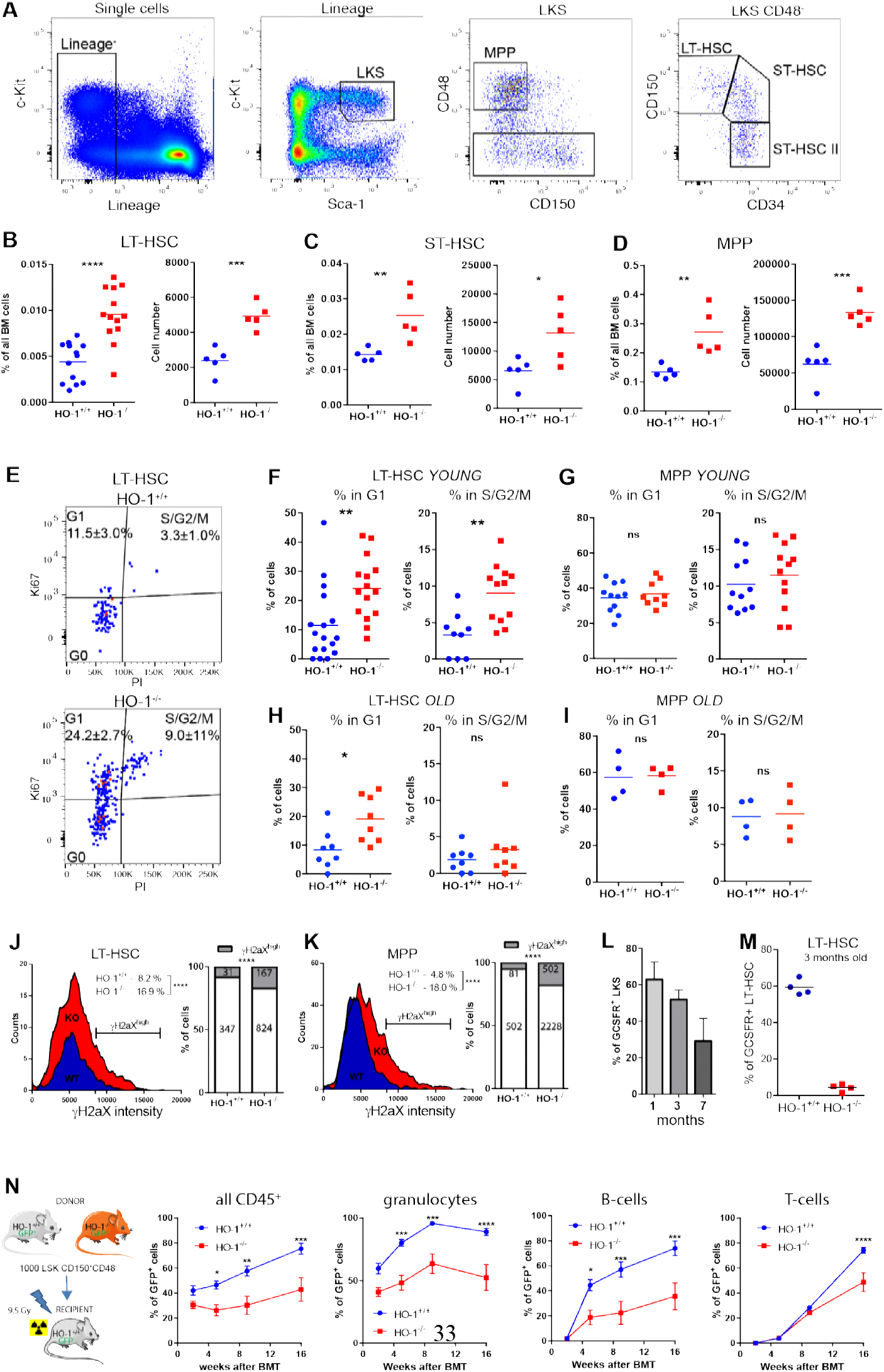
HO-1 deficiency causes loss of LT-HSC quiescence and expansion of stem cell pool. (A) Gating strategy of hematopoietic stem and progenitor cells. (B) Young HO-1^-/-^ mice possess higher number of LT-HSCs, (C) ST-HSCs and (D) MPPs. (E) Exemplary analysis of cell cycle with Ki67 and nuclear dye. (F) More young HO^-/-^LT-HSCs are in G1 and S/G2/M cell cycle phases. (G) Young HO^-/-^MPPs do not differ in cell cycling from young HO-1^+/+^ MPPs. (H) More old HO^-/-^LT-HSCs are in G1 phase in comparison to old HO-1^+/+^ LT-HSCs, but not in S/G2/M phase. (I) Old MPPs do not differ in cell cycling between genotypes. The presented cell cycle analysis is from two independent experiments. (J) Young HO^-/-^LT-HSCs contain more γH2aX^high^ cells than young HO-1^+/+^, 378-991 cells analyzed from 7 mice/group. (K) Young HO-1^-/-^MPPs contain more γH2aX^high^ cells than young HO-1^+/+^, 1693-2790 cells analyzed from 7 mice/group. (L) Percentage of GCSFR+ cells among HSPC decreases with age, n = 4/group. (M) Percentage of GCSFR+ cells among young LT-HSCs is lower in HO-1^-/-^ mice. (N) HSCs from young HO-1^-/-^ mice provide worse hematopoietic reconstitution potential after transplantation than HO-1^+/+^ HSCs, n = 8-9 mice/group

Next, we measured the levels of phosphorylated H2aX (γH2aX), an established marker of aged LT-HSCs, linked to increased DNA damage and replication stress [7,37,38]. Levels of γH2aX were significantly higher in LT-HSCs (Fig 3J) and MPPs (Figure 3K) in young HO-1^-/-^ in comparison young wildtype mice. We also noticed that in young wildtype HO-1^+/+^ mice a lower percentage of MPPs as compared to LT-HSCs is p-γH2aX^high^ (Figure 3J,K), which is consistent with recent reports indicating that DNA repair is activated upon cell cycle entry and differentiation into MPPs [39].

Importantly, we identified a novel marker of aged HSPCs. We found that a fraction of HSPCs that have G-CSF receptor on their surface (G-CSFR+) decreased with age (Figure 3L). LT-HSCs pool from young HO-1^-/-^ mice contained significantly fewer G-CSFR+ cells compared to young HO-1^+/+^ mice, similar to the reduced fraction of G-CSFR+ cells among HSPCs seen in aged HO-1^-/-^ and HO-1^+/+^ mice (Figure 3M). Interestingly, only the levels of G-CSFR protein on the cell surface were decreased, while mRNA levels were higher in aged LT-HSC (Table S5).

Finally, we assessed the functionality of HO-1^-/-^ -derived LT-HSCs in a transplantation model (Figure 3N). Transplanted HO-1^-/-^ LT-HSCs were less efficient at reconstituting irradiated HO-1^+/+^ recipients than control HO-1^+/+^ LT-HSCs in all tested lineages (granulocytes, B-cells, T-cells) (Figure 3N). In conclusion, the LT-HSCs from young HO-1^-/-^ mice exhibit features that resemble the phenotype of aged LT-HSCs, including an expanded pool, more γH2aX^high^ cells, decreased fraction of G-CSFR+ cells, and impaired regenerative potential.

### Transcriptome profiling of young HO-1^-/-^ LT-HSC reveals their premature aging

Our analysis showed that LT-HSCs isolated from young HO-1^-/-^ mice display markers of aged LT-HSCs. Therefore, we hypothesized that HO-1 deficiency may contribute to the premature aging of LT-HSCs. To test this hypothesis in the most comprehensive way possible, we profiled the transcriptome of LT-HSCs from young (3-months old) and aged (~18 month old) mice of HO-1^+/+^ and HO-1^-/-^ genotype by RNA-sequencing.

First, we compared LT-HSCs from young HO-1^-/-^ and HO-1^+/+^ mice to determine the effect of HO-1 deficiency on the LT-HSC transcriptome. We identified 1067 significant DEGs (FDR < 0.1) between LT-HSCs from young HO-1^-/-^ and young HO-1^+/+^ mice, which segregated these groups into distinct clusters (Figure 4A, Table S4). Similarly, we identified 595 significant DEGs (FDR = 0.1) between LT-HSCs from old HO-1^-/-^ and old HO-1^+/+^ mice, which segregated these groups into distinct clusters (Figure 4B, Table S4). We performed GSEA using the Gene Ontology Biological Process (GOBP) database to ask which biological processes are associated with these DEGs. The comparison of HO-1^-/-^ and HO-1^+/+^ LT-HSCs from both young and old mice identified similar GOBP terms (Figure 4C,D). We found that the significant pathways were associated with cell aging, cell cycle, and DNA damage repair, consistent with our flow cytometry results of cell cycle status and γH2aX expression (Figure 3, Figure S3). The DEGs were additionally implicated in cell adhesion, metabolic control, and regulation of hematopoietic differentiation (Figure 4C,D). Interestingly, the GSEA analysis also identified processes related to response to reactive oxygen species (ROS) and response to hypoxia, but only between old HO-1^-/-^ and HO-1^+/+^ LT-HSC (Figure 4D).

**Figure 4.**
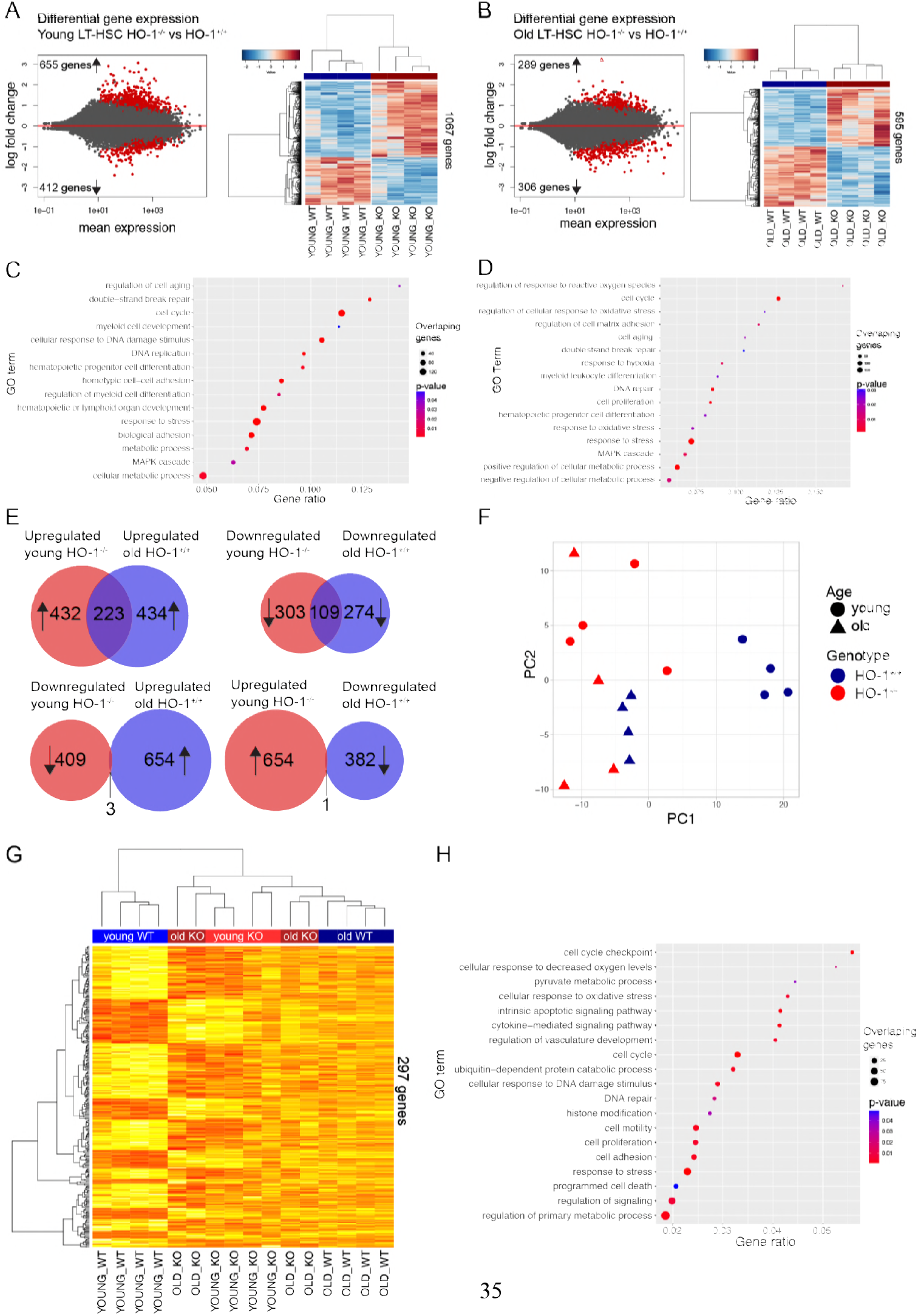
LT-HSCs from young HO-1^-/-^ mice possess transcriptional profile indicating premature aging. (A) RNA-seq revealed 1067 DEGs between young HO-1^-/-^ and HO-1^+/+^ LT-HSCs. (B) RNA-seq revealed 595 DEGs between old HO-1^-/-^ and HO-1^+/+^ LT-HSCs. (C) Selected GOBP terms enriched in GSEA analysis in young HO-1^-/-^ LT-HSCs. (D) Selected GOBP terms enriched in GSEA analysis in old HO-1^-/-^ LT-HSCs. (E) Significant part of DEGs (31% total) in young HO-1^-/-^ LT-HSC overlaps with DEGs identified in LT-HSC during physiological aging. (F) PC1 in PCA analysis on selected 1148 genes separated young HO-1^+/+^ LT-HSC from other groups, clustering the young HO-1^-/-^ LT-HSC together with old LT-HSC. (G) Hierarchical clustering on the 20% of most correlated 297 genes with PC1 showed 2 major clusters that includes young HO-1^+/+^ LT-HSC in one and old LT-HSC with young HO-1^-/-^ LT-HSCs in second. (H) Selected GOBP terms enriched in GSEA analysis based on 297 identified genes.

Next, we asked whether the genes dysregulated in young HO-1^-/-^ LT-HSCs resemble the expression changes in LT-HSCs during aging. First, we identified 1040 significant gene expression changes (FDR < 0.1) in LT-HSCs acquired with age by comparing young HO-1^+/+^ LT-HSC and old HO-1^+/+^ LT-HSC (Table S5). Then, we compared the DEGs between aged HO-1^+/+^ and young HO-1^+/+^ LT-HSCs with the DEGs between young HO-1^-/-^ and young HO-1^+/+^ LT-HSCs. We found 223 shared DEGs that are upregulated in both young HO-1^-/-^ LT-HSCs and old HO-1^+/+^ LT-HSCs compared to young HO-1^+/+^ LT-HSCs (Figure 4E, Table S6), and 109 shared DEGs that are downregulated in young HO-1^-/-^ LT-HSCs and old HO-1^+/+^ LT-HSCs compared to young HO-1^+/+^ LT-HSCs (Figure 4E, Table S6). In contrast, we found only 4 DEGs that are significantly changed in opposite directions in young HO-1^-/-^ LT-HSCs and old HO-1^+/+^ LT-HSC (Figure 4E). This suggests that 31% of DEGs between young HO-1^-/-^ LT-HSCs and young HO-1^+/+^ LT-HSCs represents HO-1-induced premature aging and are shared with the DEGs found between aged HO-1^+/+^ LT-HSC and young HO-1^+/+^ LT-HSC, which represents normal aging.

To identify the core genes contributing to premature aging by HO-1 deficiency, we asked whether there are genes similarly expressed in young HO-1^-/-^ LT-HSCs and old LT-HSCs regardless of the genotype, but differentially expressed in young HO-1^+/+^ LT-HSCs. We found 1448 DEGs that satisfied these criteria and performed principal component analysis (PCA) based on the identified DEGs. The PC1 component separated the young HO-1^+/+^ LT-HSCs from the other samples and grouped young HO-1^-/-^ LT-HSC with old HO-1^-/-^ and HO-1^+/+^ LT-HSC (Figure 4F). To identify the most significant genes that provide this separation, we calculated the top 20% of most correlated (both positively and negatively) genes with PC1. We identified 297 genes that clustered young HO-1^+/+^ LT-HSCs separately and young HO-1^-/-^ LT-HSCs together with old HO-1^-/-^ and old HO-1^+/+^ LT-HSCs (Figure 4G). GSEA revealed that these 297 genes are involved in a broad variety of biological processes, including cell cycle, DNA repair, metabolism, cell adhesion, response to stress, signaling, histone modification, protein catabolic processes, and apoptosis (Figure 4H). Particularly, this GSEA analysis above indicated that alterations in expression of genes regulating pyruvate metabolism converge the young HO-1^-/-^ LT-HSCs with old LT-HSCs. Pyruvate metabolism and the regulation of glycolysis are quiescence checkpoints in HSCs [40]. Decreased expression of genes regulating pyruvate and glycolysis in LT-HSCs, like pyruvate dehydrogenase kinases (*Pdks*) and lactate dehydrogenases (*Ldhs*), respectively, is linked with lower ATP levels [40]. We found that young HO-1^-/-^ LT-HSCs downregulate expression of *Ldhb, Pdk2*, and *Pdpr*, similarly to old LT-HSCs (Figure 5A). Consistently, we observed that HO-1 deficiency results in lower amounts of ATP in stem and progenitor cells in young mice (Figure 5B) but not in old mice (Figure 5C).

**Figure 5.**
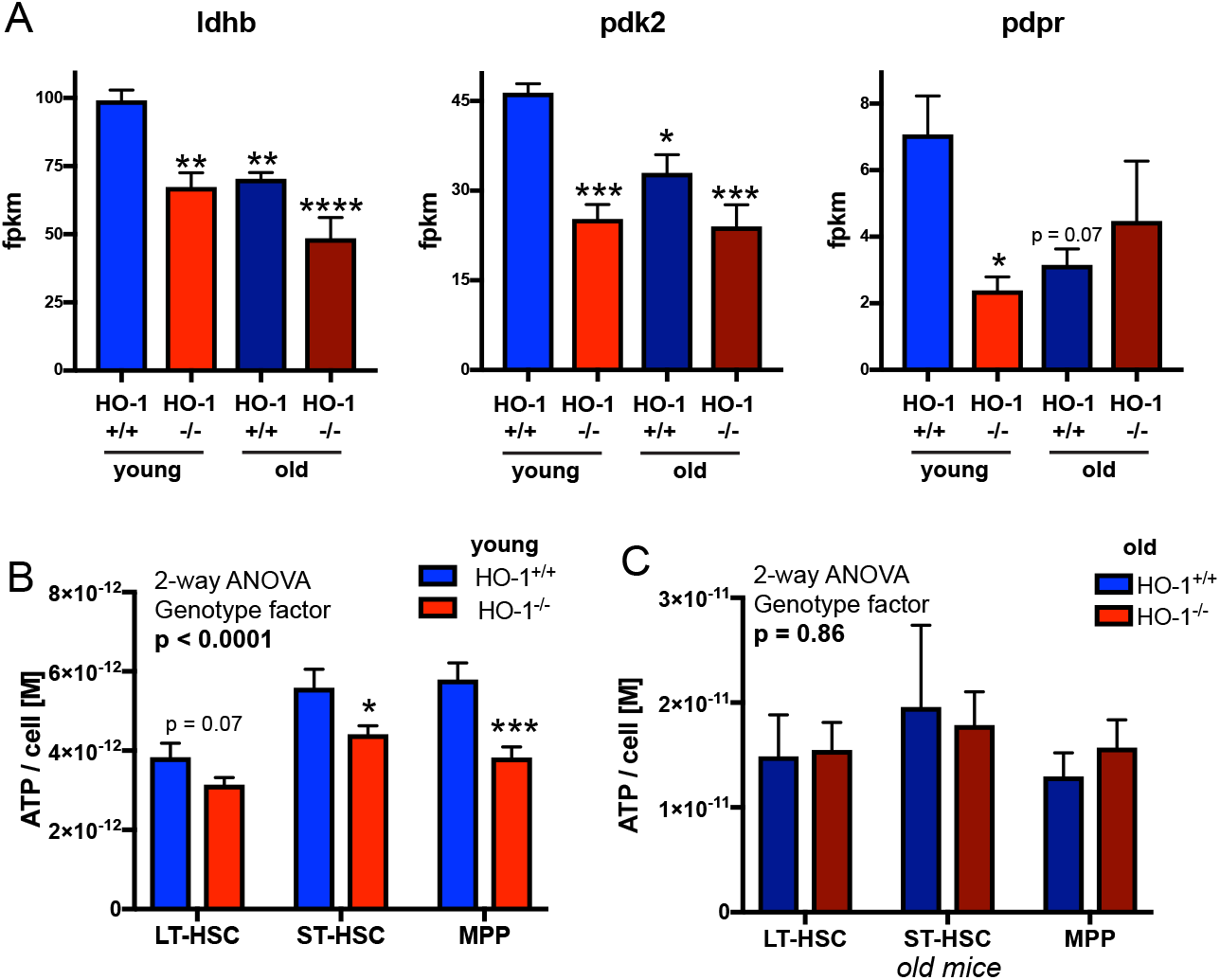
Decreased expression of genes regulating pyruvate metabolism in young HO-1^-/-^ LT-HSCs and old LT-HSCs is linked with lower ATP levels. (A) Ldhb, Pdk2 and Pdpr are downreglated in young HO-1^-/-^ LT-HSCs, but not in old young HO-1^-/-^ LT-HSCs. Analyzed by RNA-seq, 4 mice/group, * - p<0.05 comparing to young HO-1^+/+^ group. (B) ATP levels are lower in young HO-1^-/-^ LT-HSCs comparing to young HO-1^+/+^ LT-HSCs, (C) but not in old HO-1^-/-^ LT-HSCs comparing to old HO-1^+/+^ LT-HSCs. ATP levels measured in two independent experiments, n = 8-18/group.

Overall, transcriptomic analysis of young HO-1^-/-^ LT-HSCs revealed similarity to transcriptome profile of old HO-1^+/+^ LT-HSC, indicating premature aging of LT-HSCs in young HO-1^-/-^ mice. We found, that transcriptome similarity of young HO-1^-/-^ LT-HSC to aged LT-HSC is manifested by expression of genes involved in variety of biological process, including downregulation of genes involved in the pyruvate metabolism, that was linked with lower amount of ATP.

### HO-1 deficiency in the niche causes exhaustion of LT-HSCs

We found that LT-HSCs from HO-1^-/-^ mice are impaired and exhibit the features of premature aging. As we demonstrated, expression of HO-1 is low in LT-HSCs, but high in macrophages, ECs and CAR cells (Figure 1E-H). Therefore, we hypothesized that the observed LT-HSCs phenotype in HO-1^-/-^ mice is caused by HO-1-deficiency in the HSC microenvironment. To verify this, we first asked whether HO-1 expression in macrophages is necessary to maintain quiescence of LT-HSCs. We analyzed the LT-HSCs in mice with conditional deletion of HO-1 with Cre recombinase driven by the lysozyme M promoter (HO-1^fl/fl^;LysM-Cre) [41]. Deletion of HO-1 by LysM-Cre in monocyte-macrophage and neutrophil lineages [41,42] did not reproduce the impaired LT-HSC phenotype observed in global HO-1-deficient mice. The HO-1^flfl^;LysM-Cre mice did not show expansion of LT-HSCs or HSPCs, and there were no differences in the cell cycle, suggesting that lack of HO-1 in monocyte-macrophage lineage and neutrophils alone do not affect LT-HSC (Figure 6A).

**Figure 6.**
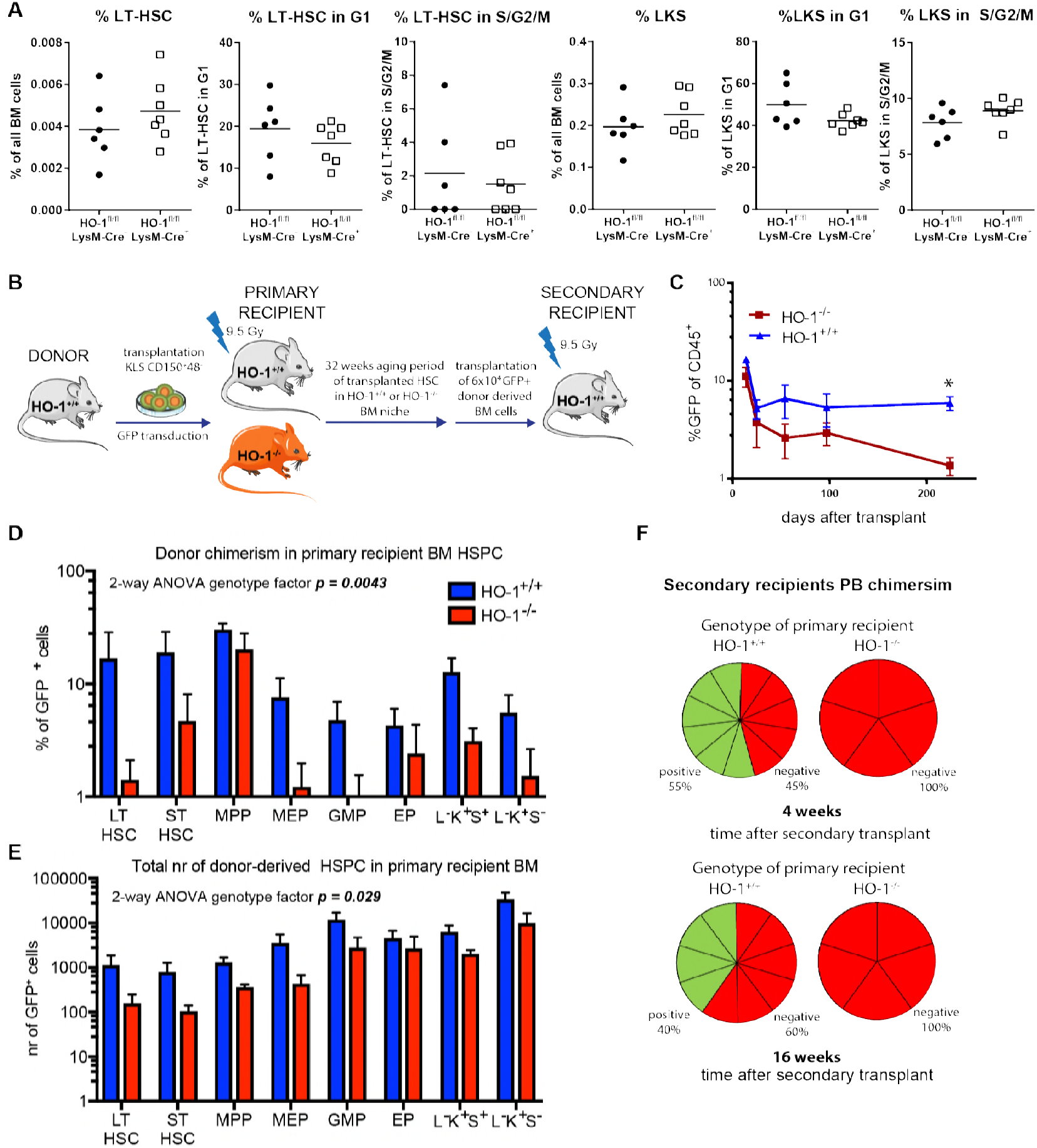
HO-1 deficiency in the niche cause exhaustion of LT-HSCs. (A) Deletion of HO-1 in myeloid lineage did not cause expansion of LT-HSCs and did not alter their cell cycle status. (B) Scheme of the experiment assessing long-term effect of HO-1 deficient niche on function of HSCs. (C) Long-term PB chimerism derived from HSCs transplanted to HO-1^-/-^ recipients is lower than from HSCs transplanted to HO-1^+/+^ recipients. (D) Chimerism among BM HSPCs fractions and (E) total number of cells derived from HSCs transplanted to HO-1^-/-^ recipients are lower than from HSCs transplanted to HO-1^+/+^ recipients. (F) HSCs that were initially transplanted to primary HO-1^-/-^ recipients did not reconstitute secondary HO-1^+/+^ recipients, in contrast to HSCs that were initially transplanted to HO-1^+/+^ recipients, n = 312/group.

Next, to investigate the role of HO-1-expressing niche cells, we transplanted HO-1^+/+^ HSCs to lethally irradiated HO-1^-/-^ or HO-1^+/+^ recipients and followed donor chimerism for 32 weeks (Figure 6B). We did not observe any increased mortality or morbidity of HO-1^-/-^ mice upon irradiation. During the first 3 weeks after transplant, we did not find any significant differences in donor chimerism between HO-1^-/-^ and HO-1^+/+^ recipients as measured by percent of GFP+CD45+ peripheral blood (PB) cells (Figure 6C). However, starting from 7 weeks after transplant, chimerism in HO-1^-/-^ recipients decreased, reaching the highest and most significant difference after 32 weeks (Figure 6C).

Looking within the bone marrow after 32 weeks, we observed lower percent chimerism (Figure 6D) and a total number of donor-derived cells (Figure 6E) in BM HSC and HSPC fractions in HO-1^-/-^ than in HO-1^+/+^ recipients. To evaluate the function of donor-derived HO-1^+/+^ HSCs that were aged for 32 weeks in HO-1^-/-^ or HO-1^+/+^ primary recipients, we serially transplanted the same number (6 x 10^4^) of donor-derived GFP+ whole BM cells to secondary young HO-1^+/+^ recipients (Figure 6B). In this experimental setting, we transplanted BM HO-1^+/+^ cells to the HO-1^+/+^ secondary recipients, and the only thing that differentiated the groups was the 32 weeks aging period in HO-1^+/+^ or HO-1^-/-^ primary recipients (Figure 6B). We found that only BM cells from HO-1^+/+^ recipients reconstituted secondary recipients, while BM cells from HO-1^-/-^ primary recipients failed to reconstitute secondary recipients (Figure 6F). These results demonstrate that HO-1 deficiency in the BM environment causes exhaustion of LT-HSC and impairs their regenerative potential.

### HO-1^+/+^ niche can restore the function and transcriptional profile of HO-1^-/-^ LT-HSC

Given that HO-1 in the niche is necessary for proper HSC function, we asked whether the HO-1^+/+^ niche can restore the function of LT-HSCs isolated from young HO-1^-/-^ mice (Figure 7A). As shown in Figure 3N, young HO-1^-/-^ HSC transplanted into primary HO-1^+/+^ recipients are less efficient at reconstituting hematopoiesis than young HO-1^+/+^ HSCs, even after 16 weeks of transplant. This indicates that primary transplantation to the HO-1^+/+^ niche is not sufficient to restore the function of young HO-1^-/-^ LT-HSCs.

**Figure 7.**
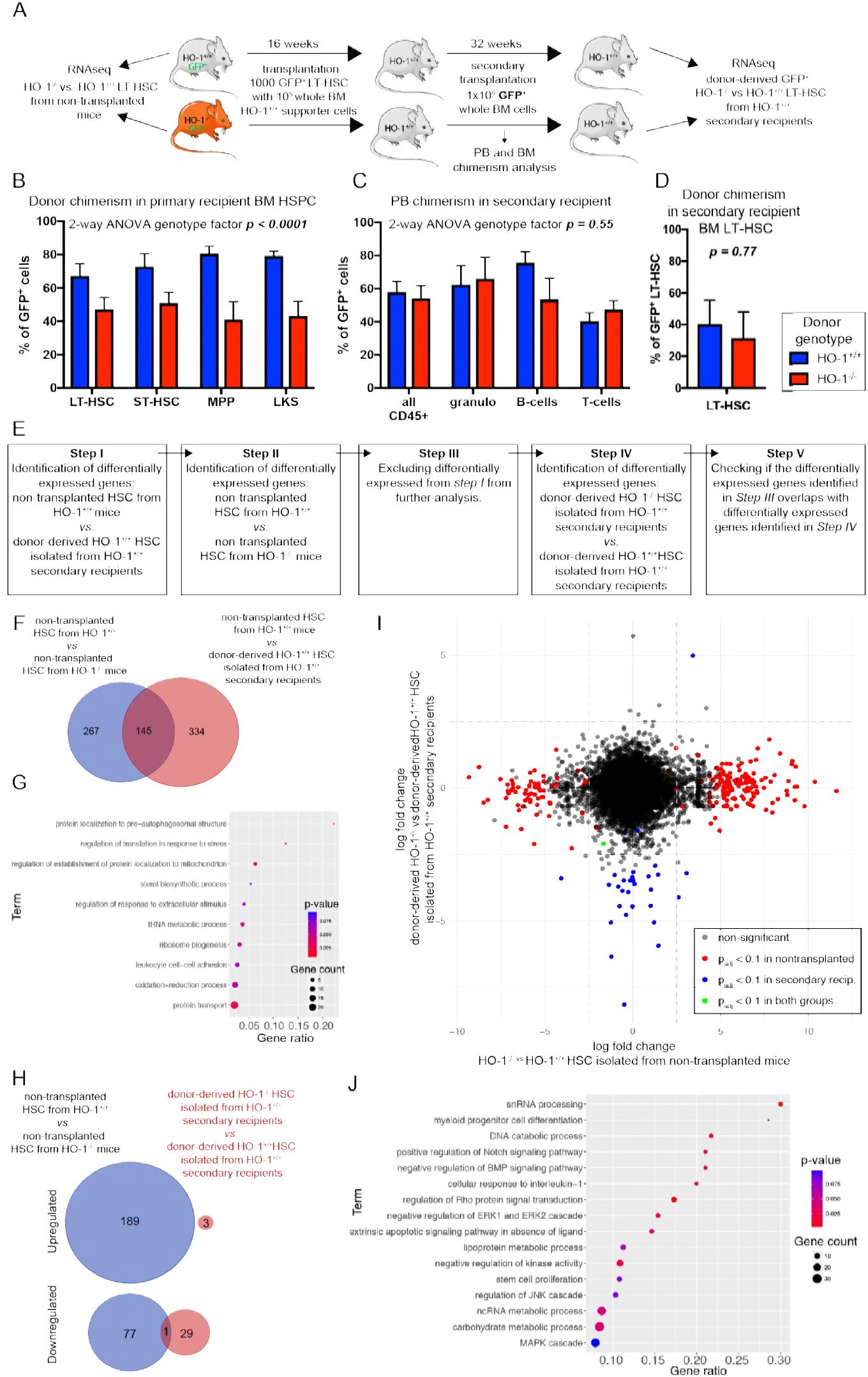
HO-1^+/+^ restore function and transcriptional profile of HO-1^-/-^ LT-HSCs. (A) Scheme of the experiment verifying if the HO-1^+/+^ niche is able to reverse phenotype of HO-1^-/-^ LT-HSCs. (B) Transplantation of HO-1^-/-^ HSCs provides lower chimerism among HSPC fractions in primary recipients, n = 7-8 mice/group. (C) Transplantation of the same number of donor-derived BM cells from primary recipients provide the same PB chimerism and (D) BM LT-HSC chimerism, n = 7-8 mice/group. (E) Analysis pipeline used to determine whether transplantation of HO-1^-/-^ HSCs to the HO-1^+/+^ recipients reverses their transcriptional alterations. (F) Overlap between DEGs in young HO-1^-/-^ HSCs and DEGs changed by transplantation alone. 145 overlapping genes were excluded from further analysis. (G) GSEA analysis based on 145 excluded genes indicates processes that cannot be analyzed with the pipeline. (H) Only 1 out of 267 DEGs identified in non-transplanted HO-1^-/-^ LT-HSCs was still dysregulated in HO-1^-/-^ LT-HSCs transplanted twice to the wild-type HO-1^+/+^. (I) Comparison of gene log fold changes in non-transplanted HO-1^-/-^ LT-HSCs and HO-1^-/-^ LT-HSCs transplanted twice to the wild-type HO-1^+/+^showed that transplantation of HO-1^-/-^ LT-HSCs twice to the wild type niche reverses their transcriptional alterations. (J) GSEA analysis based on genes that were altered in non-transplanted HO-1^-/-^ LT-HSCs, but were normalized by double transplantation to the wild-type niche.

However, primary recipients transplanted with HO-1^-/-^ LT-HSCs have a lower donor-derived chimerism among HSPCs (Figure 7B), which may explain the lower PB chimerism. To test whether secondary transplantation into the HO-1^+/+^ niche can restore the function of HO-1^-/-^ LT-HSCs, 16 weeks after primary transplant we transplanted equal numbers of GFP+ donor-derived BM cells from HO-1^+/+^ primary recipients containing GFP+HO-1^+/+^ chimeric BM or GFP+HO-1^-/-^ chimeric BM into HO-1^+/+^ secondary recipients (Figure 7A). The frequency of LT-HSCs in donor-derived BM cells was the same in both groups (Figure S4). After secondary transplantation of LT-HSCs initially derived from HO-1^-/-^ and HO-1^+/+^ donors, recipient mice exhibited no difference in PB chimerism in any of tested lineages (Figure 7C), and no difference in chimerism among BM LT-HSCs 36 weeks later (Figure 7D). This indicates that the HO-1^+/+^ niche can restore function of HO-1^-/-^ LT-HSCs after secondary transplantation to HO-1^+/+^ recipients.

Additionally, we compared the transcriptome of freshly isolated LT-HSCs from HO-1^+/+^ and HO-1^-/-^ mice, as well as from donor-derived HO-1^+/+^ and HO-1^-/-^ LT-HSCs isolated from secondary recipients 36 weeks after transplantation (Figure 7A). By sorting similar numbers of identically defined LT-HSCs and processing all samples with the same RNA-seq library preparation and sequencing steps, we verified whether the secondary HO-1^+/+^ niche reverses the defective transcriptome of LT-HSCs freshly isolated from HO-1^-/-^ mice. Given that transplantation alone changes the transcriptome of LT-HSCs, we identified and then excluded the genes whose expression significantly changes with transplantation (Figure 7E). To identify genes that are affected by transplantation alone, we compared freshly isolated, non-transplanted HO-1^+/+^ LT-HSCs with serially transplanted HO-1^+/+^ LT-HSC isolated from the secondary recipient. We identified 479 DEGs (Figure 7E step I) and subsequently excluded them from our analysis. Next, we identified 412 DEGs between freshly isolated, non-transplanted LT-HSCs from HO-1^+/+^ and HO-1^-/-^ mice, of which 267 DEGs did not overlap with previously excluded genes that were changed by transplantation alone (Figure 7E, step II, III, Figure 7F) and were included in the next steps of analysis. 145 overlapping DEGs that were excluded from the analysis at this step (Figure 7F, genes there were changed between non-transplanted HO-1^+/+^ and HO-1^-/-^ LT-HSCs and changed by transplantation alone) were used for GSEA to indicate processes that cannot be addressed with this experimental scheme (Figure 7G).

Finally, we checked whether secondary transplantation to wildtype HO-1^+/+^ recipients reverses transcriptional alterations of HO-1^-/-^ LT-HSCs. When we compared HO-1^+/+^ and HO-1^-/-^ LT-HSCs from secondary recipients, we identified only 33 DEGs (Figure 7G, step IV). Next, we verified how many of 267 DEGs identified in non-transplanted HO-1^-/-^ LT-HSCs are still dysregulated in HO-1^-/-^ LT-HSCs transplanted twice to the wild-type HO-1^+/+^ secondary recipients (Figure 7E, step V), and we found only 1 overlapping gene (Figure 7H).

We also compared fold changes of DEGs identified between non-transplanted HO-1^-/-^ LT-HSCs and HO-1^+/+^ LT-HSCs to fold changes of DEGs identified between serially transplanted HO-1^-/-^ LT-HSCs and HO-1^+/+^ LT-HSCs from the secondary recipients (Figure 7I). The observed pattern indicates that most genes dysregulated in non-transplanted HO-1^-/-^ LT-HSCs were normalized in HO-1^-/-^ LT-HSCs that were transplanted twice into HO-1^+/+^ recipients (Figure 7I). The GSEA based on genes that were normalized upon transplantations (2.5 log-fold changed in young in nontransplanted HO-1^-/-^ LT-HSC, but less than 2.5 log-fold changed in transplanted HO-1^-/-^ LT-HSCs from the secondary recipients, Figure 7I) implicated processes related to snRNA processing and ncRNA metabolism, myeloid differentiation, Notch signaling pathway, BMP signaling pathway, Rho signaling, several metabolic processes, as well as processes and pathways involved in proliferation (Figure 7J). In summary, serial transplantation of HO-1^-/-^ LT-HSCs into HO-1^+/+^ recipients restores function and reverses transcriptional alterations of HO-1^-/-^ LT-HSCs.

## Discussion

We demonstrated that HO-1 is highly expressed in the BM niche. HO-1 deficiency affects the BM niche and is linked to premature exhaustion of HSCs. Expression of HO-1 in the niche decreases with age. In accordance, LT-HSCs in HO-1-deficient mice prematurely acquire aging phenotype, as revealed by their impaired function and aged-like transcriptional signature. Notably, the impaired function and alterations of HO-1^-/-^ LT-HSCs are reversed by serial transplantation into young wildtype HO-1^+/+^ recipients.

In this study, we characterize the aging of HSCs not only by known hallmarks of aging but also by global transcriptional profiling (Figure 4). The classical hallmarks of HSC aging include expansion, increased markers of DNA damage, myeloid-biased differentiation, loss of protein polarity, and reduced reconstitution capacity [3,4,7,1,43]. However, these features may not be specific to the physiological process of aging and can be just derivatives of other dysregulations, such as altered cell cycle control or impaired DNA repair mechanism. Therefore, we profiled the transcriptome of young and aged, wildtype and mutant HO-1 LT-HSCs within the same sequencing and analysis procedure as a new and comprehensive method to evaluate the aging phenotype. Using this approach, we found transcriptional similarity between LT-HSCs isolated from young HO-1^-/-^ mice and LT-HSCs isolated from physiologically aged wild-type mice of matching genotypic strains. This similarity pertains to genes involved in broad kinds of processes: cell cycle, metabolism, cell adhesion, DNA repair, and cytokine signaling (Figure 4H). The altered expression of cell cycle genes in old LT-HSCs is consistent with previous work on the transcriptome of aged HSCs [44], although we did not observe an increase in the percentage of LT-HSCs in the G1 phase in old wildtype mice, as reported previously [44]. Nevertheless, we observed that LT-HSCs from young HO-1^-/-^ mice have an increased fraction of cells in G1 and S/G2/M phases compared to young wildtype HO-1^+/+^ (Figure 3F), while LT-HSCs from older HO-1^-/-^ mice have more cells in G1 phase compared to older HO-1^+/+^ mice, but not in S/G2/M phases (Figure 3H). The latter results can be explained by generally decreased proliferation capability of old HSCs as shown previously [2].

The other characteristic of aging LT-HSCs is the dysregulation of metabolic checkpoints. Sirt-7 has been proposed as one of the metabolic regulators that are decreased in aged LT-HSC [45]. Consistently, we observed significantly reduced expression of Sirt-7 in aged wild-type LT-HSCs (Table S5) and similar trends in young HO-1^-/-^ LT-HSCs (Table S3). We found that genes regulating pyruvate metabolism and maintaining glycolysis in HSCs are also decreased in old LT-HSCs and young HO-1^-/-^ LT-HSCs (Figure 5A). Downregulation of these genes in young HO-1^-/-^ mice is linked to reduced levels of ATP in HSPCs (Figure 5B). This is consistent with previous work demonstrating that HIF-1α^Δ/Δ^ LT-HSCs with decreased Pdk2 and Pdk4 expression also have reduced ATP levels [40].

In the present work, we did not use myeloid skewing as a marker of LT-HSC aging. Although, we do observe significantly more myeloid cells in PB of HO-1^-/-^ mice (data not shown), we previously showed that the myeloid bias in HO-1^-/-^ mice is linked to HO-1’s role at the level of myelocytes [46]. Given that HO-1-deficiency causes the myeloid bias at the level of myelocytes, aging of LT-HSC in HO-1^-/-^ animals could not be judged by increased output of mature myeloid cells.

Finally, while we observed the broad similarities of transcriptome of HSCs from young HO-1^-/-^ mice to HSC from old physiologically aged HO-1^+/+^ mice, some features differ between these two groups. It is known that in old mice HSC fractions expands but the MPPs fraction does not [47]. In contrast, in young HO-1^-/-^ animals we observed expansion of both HSCs and MPPs (Figure 3B,D). Therefore, while whole transcriptional profile and phenotype of HSCs isolated from young HO-1^-/-^ animals generally resemble aged HSCs, there are some features that differ HSCs from HO-1^-/-^ animals from old HSCs.

Two previous papers assessed the role of HO-1 in HSCs, however they did not focus on steady state function of the niche. Cao and colleagues showed that HO-1+^-^ mice had normal hematopoiesis in steady-state conditions, but showed impaired long-term hematopoietic activity upon 5-FU induced myelotoxic injury and serial transplantation [48]. Other group revealed that HO-1 inhibition may be strategy to improve mobilization of HSC [49]. While, the conclusions of these papers are in line with our work, here we describe a broader role of HO-1 on strictly defined HSC. For the first time, we showed that HO-1 should not only be considered as intrinsic enzyme activated in various hematopoietic cells in stress conditions, but also as crucial extrinsic regulator of strictly defined HSCs in steady state conditions.

Our transcriptional data indicating the premature aging of HO-1^-/-^ LT-HSCs have been supported by functional *in vivo* studies. We have proposed novel schemes of HSC transplantation to test the role of the investigated niche-factor in HSC aging (Figure 6B). Namely, we transplanted HSCs from wildtype mice into recipients deficient in HO-1 to demonstrate niche-mediated changes, and then performed secondary transplant of the same number of only donor-derived BM cells to young wild-type animals. This allowed us to prospectively check how aging of LT-HSCs in HO-1^-/-^ mice affects their function and to demonstrate niche-mediated reversal of this premature aging. We believe that our experimental strategy may be used to study the role of other niche factors.

We did not observe any increased morbidity or mortality of HO-1^-/-^ mice upon irradiation. All mice survived the conditioning, demonstrating that HO-1^-/-^ mice can be used in standard transplantation protocols as recipients. Furthermore, we did not observe any short-term effects on hematopoietic parameters in HO-1^-/-^ recipients (Figure 6C), indicating that there are no differential effects of irradiation on these mice. Oppositely, we observed long-term effects on hematopoiesis, especially during secondary transplantation assay (Figure 6C,F), what is thought to be classical output to measure HSC function [50].

While HSC aging is a known phenomenon, the reversal of age-related loss of HSC function has remained elusive. Here, we showed that the premature phenotype of LT-HSCs from young HO-1^-/-^ LT-HSCs can be reversed by the wildtype niche. Upon primary transplantation into wildtype recipients, HO-1^-/-^ LT-HSCs demonstrated impaired reconstitution even after 16 weeks, without any signs of a reversal of declined function (Figure 3N). However, as we showed, LT-HSCs isolated from young HO-1^-/-^ mice have altered gene expression profiles (Figure 4) and cell cycle status (Figure 3F). Therefore, it is possible that they lost the competition for niches with residual wildtype LT-HSCs and LT-HSCs within the supporter cells at the initial stages after transplantation, and they are not able to rebound the contribution to blood production afterwards. Indeed, we observed lower chimerism among the HSPC fraction after transplantation of HO-1^-/-^ HSCs (Figure 6D). Finally, while HO-1 is not expressed in HSCs in steady-state it might be expressed upon stress conditions [48]. Thus, lack of HO-1 during the transplantation stress may affect their viability. To avoid this equivocally interpretations we did serial transplantion of the same number of donor-derived cells from primary recipients into the secondary recipients. By this way we provided the same conditions for the compared groups. In such experimental settings, the secondary HO-1^+/+^ niche fully restored the reconstitution potential of HO-1^-/-^ HSCs to equal that of control HO-1^+/+^ HSCs (Figure 7C). This implies that the young HO-1^+/+^ niche can reverse the impaired reconstitution capacity of prematurely aged HO-1^-/-^ HSCs and that the induction of HO-1 in HSCs during transplantation is not limiting factor in the assay.

A recent study demonstrated that reduced expression of osteopontin (*Spp1*) in BM stroma regulates aging of HSCs and restoring Spp1 levels attenuates aged phenotype of HSCs [51]. This is in line with our study indicating that the young niche has the potential to reverse some of the aging-related features of HSC. However, we did not observe altered expression of *Spp1* in HO-1^-/-^ ECs or CARs (Table S1, S2). Instead, we suppose that *Scf* and *Sdf1* possibly mediate premature aging of HSC in case of HO-1 deficiency. *Scf* and *Sdf1* role in the maintenance of HSC is well evidenced [25,26] and we did observe downregulation of these factors in HO-1^-/-^ niche populations, with the strongest decrease of *Scf* in CD31^+^Sca-1^+^ EC. Interestingly, young HO-1^-/-^ LT-HSC and old HO-1^+/+^LT-HSC showed higher expression of *Cxcr4* – a receptor for Sdf1 – what might indicate a compensatory mechanism of decreased levels of *Sdf1.*

It was recently proposed that a particular fraction of endothelial cells called H-type is crucial for HSC maintenance and regulates their aging. The fraction of H-type ECs produces high levels of Scf, but their numbers decrease with age [52]. This is also consistent with our data on lower CD31^+^Sca-1^+^ EC numbers in old animals and significantly reduced expression of *Scf* by HO-1^-/-^ CD31^+^Sca-1^+^ ECs (Figure 1J, Figure 2E, Table S1), although we did not specifically distinguish H-type EC cells in our experiments.

Interestingly, the decreased production of *Scf* and *Sdf1* by HO-1^-/-^ niche is linked with different HSC phenotype than total genetic ablation of *Scf* or *Sdf1* in niche cells. We observed that reduced levels of *Scf* and *Sdf1* relates to activation and expansion of HSC, while total deletion resulted in collapse of HSCs or mobilization of HSC into peripheral blood [26,25,27,22].

Along with demonstrating that function of HO-1^-/-^ LT-HSCs can be restored by HO-1^+/+^ niche, we checked whether the transcriptional profile of HO-1^-/-^ LT-HSC was also reversed. Naturally, this comparison has limitations, as transplanting the cells twice alone induces their aging [53] as well the duration of experiment can cause significant transcriptional changes. Thus, it is not unreasonable that a large number of genes that were changed in LT-HSCs from serial transplantation overlap with genes that are differentially expressed between the impaired young HO-1^-/-^ LT-HSCs and the wildtype young HO-1^+/+^ LT-HSCs (Figure 7F,G). Although these overlapping genes had to be excluded from our analysis given the confounding effect of transplantation itself, we still demonstrated that transplantation of HO-1^-/-^ LT-HSCs into the HO-1^+/+^ niche reverses the altered expression of the other non-overlapping, differentially expressed genes (Figure 7I).

Our findings suggest that reversing the age-related decline in HSC function may be achieved by enhancing the HO-1 activity in the niche. Potential clinical translation of HO-1 function in the HSC niche should take into consideration: 1) monitoring HO-1 expression as a diagnostic for bone marrow age and niche dysfunction, 2) screening for HO-1 polymorphisms in hematological disorders, 3) targeting HO-1 activity and expression by pharmacological induction. In humans, the activity of HO-1 is linked to promoter polymorphisms [54,55]. Longer promoters with more GT repeats are linked to lower expression of HO-1, while shorter GT repeats lead to higher levels of HO-1 [55,34]. We showed previously that in human endothelial cells the shorter alleles of HO-1 promoter provide better response to oxidative stress, VEGF-induced proliferation, and lower production of inflammatory cytokines [34]. Given the crucial role of endothelial cells in the HSC niche, people with longer HO-1 promoter variant and impaired function of endothelial cells might have increased risk of accelerated aging of the HSC. Secondly, some kinds of porphyrins, e.g., cobalt protoporphyrin IX, induce expression of HO-1 *in vivo* and might be considered as potential treatment in case the function of HSC is impaired. Taken together, this study suggests an important role of HO-1 also in the human HSC niche and calls for future studies to pursue the clinical applications of our findings.

We demonstrated high HO-1 expression in steady state conditions in BM niche. Therefore, we focused on the HO-1 role in the BM niche. We characterized which populations express HO-1, how the expression changes in aging, and demonstrated that lack of HO-1 in niche cells dysregulates production of hematopoietic fractions. Given our findings on role of HO-1 in BM niche, it is likely that lack of HO-1 in the niche cells drives premature exhaustion of HSCs in HO-1^-/-^ mice. However, we cannot exclude that other systemic factors intertwine with HO-1 role within the BM niche and together contribute to dysregulation of HSC in HO-1^-/-^ mice. Investigating effects of HO-1 deficiency in BM niche separately from potential systemic effects of HO-1 deficiency using conditional genetic models (eg. VeCad-Cre, Lepr-Cre, Prx1-Cre) would be also equivocal, given that HO-1 is expressed in several niche populations (ECs, CARs, PaSs). Specific deletion of HO-1 in all niche populations using conditional genetic models would be technically complex.

While our HO-1^-/-^ mice represents the constitutive knock-out model the observed phenotype of HSC in BM of young mice is not likely caused by developmental alterations and colonization of BM by HSCs, as existing data indicates no HO-1 expression in fetal liver HSCs [56].

In conclusion, we identified HO-1 as a new factor in the BM niche. The expression of HO-1 in the BM niche decreases in aged animals and extrinsic HO-1 deficiency is linked to premature exhaustion of LT-HSCs, as shown by the loss of function and global transcriptional profiling. Notably, the premature aging phenotype of LT-HSC in the HO-1-deficient niche can be reversed by transplantation to young HO-1^+/+^ niche. This study suggests that modulation of HO-1 activity in BM niche may be used to restore the impaired function of aged HSCs.

## Acknowledgments

We would like to acknowledge Ewa Werner, Karolina Hajduk and Janusz Drebot for their help and support in performing mouse experiments as well as Agnieszka Andrychowicz-Róg and Joanna Uchto for technical support. The study was supported by the National Science Center (grant Harmonia no NCN2015/18/M/NZ3/00387 awarded to A.J. and grant Preludium no NCN2013/11/N/NZ3/00956 awarded to K.S.) and by structural funds from the EU (grant 01.02.00-069/09). A.S. and K.S. are supported by Mobility Plus grants financed by the Ministry of Science and Higher Education. The Faculty of Biochemistry, Biophysics, and Biotechnology of Jagiellonian University is a partner of the Leading National Research Center (KNOW) supported by the Ministry of Science and Higher Education

## Author Contributions

Conceptualization: KS, AJ, JD. Formal Analysis: KS, MZ, GG. Investigation: KS, MZ, AS, WN, MC, KBS, NKT, AK, EE, JK, SC. Methodology: EE, HE. Writing: Original Draft KS, AJ, AS, GG, IW. Supervision: AJ, JD.

## Methods

### Animals

All animal procedures and experiments were performed in accordance with national and European legislations, after approval by the First Local Ethical Committee on Animal Testing at the Jagiellonian University in Krakow (approval number: 56/2009, 113/2014).

The HO-1^-/-^ mice and HO-1^-/-^ mice constitutively expressing GFP (HO-1^-/-^GFP+) were kindly provided by Dr. Anupam Agarwal from University of Alabama at Birmingham. HO-1^-/-^ strain poorly breed on pure C57/Bl6 background (5.1 % of expected HO-1^-/-^ pups) and therefore is maintained on mixed C57/Bl6 x FVB background (20.1% of expected HO-1^-/-^ pups, when HO-1^-/-^ males are crossed with HO-1+^/-^ females) [57]. While there is increased perinatal mortality of HO-1^-/-^ animals, we did not observed decreased lifespan among HO-1^-/-^ mice that were born live. The weight of HO-1^-/-^ mice during aging in steady-state conditions did not differ and HO-1^-/-^ mice did not reveal acute sickness or abnormalities.

To assure relative homogeneous background, all HO-1^+/+^ controls were C57/Bl6xFVB littermates from the same breeders used to obtain HO-1^-/-^ mice. HO-1^fl/fl^;LysM-Cre were analyzed at Center of Translational Research, Medical University of Vienna, Austria.

### Flow cytometry analysis and cell sorting

Flow cytometry analysis were done on LSR II and LSR Fortessa cytometers (BD Sciences). Cell sorting was done on MoFlo XDP cell sorter (Beckman Coulter). The populations used in studies were defined as follows: HSCs – LKS CD150+CD48^-^, LT-HSCs – LKS CD150+CD48^-^CD34^-^, ST-HSCs – LKS CD150+^/mid^CD48^-^CD34+, ST-HSCs II – LKS CD150^-^CD48^-^CD34+, MPP – LKS CD150^-^CD48+, ECs – CD45^-^Ter119^-^PDGFRα^-^CD31^+^Sca-1^+^, CARs – CD45^-^Ter119^-^ PDGFRa+CD31^-^Sca-1^-^, PaS – CD45^-^Ter119^-^PDGFRα+CD31^-^Sca-1+. Following antibody clones were used in the study: CD34 (clone RAM34, BD Biosciences), Sca-1 (D7, BD Biosciences, eBioscience), c-Kit (clone 2B8, eBioscience, USA), CD45 (clone 30F-11, BD Biosciences), CD150 (clone TC15-12F12.2, Biolegend), CD48 (clone HM-48-1, Biolegend), CD45R-PE (clone RA3-6B2, BD Biosciences), Ly6G and Ly6C-PE (clone RB6-8C5, BD Biosciences), TCRγδ-PE (clone GL3, BD Biosciences), TCRβ-PE (clone H57-597, BD Biosciences), CD11b-PE (clone M1/70, BD Biosciences), Ter119-PE (clone TER119, BD Biosciences), CD3 (clone 17A2, BD Biosciences), F4/80 (clone BM8, eBioscience), MHCII (clone M5/114.15.2), CD31 (clone MEC13.3, BD Biosciences), CD140α (clone APA5, eBioscience), Ki67 (clone B56), HO-1 (SPA894, Enzo Life Sciences). Intracellular staining for flow cytometry analysis were done using IntraSure Kit (BD Biosciences). Analysis of the cell cycle and measurement of the ATP content was done according to the protocols described in detail previously [58,59].

### Immunohistochemistry

Tibias and femurs were fixed in 4% PFA with 10% EDTA for 4-8 hours on ice in 4^0^C while rotating, followed by overnight incubation in 20% EDTA in 4^0^C. Next, the bones were put to sucrose solutions in PBS: 10% – 2 h, 20% – 2 h and 30% overnight in 4^0^C. Next, the excess of sucrose was removed, bones were embedded in Tissue-Tek Oct media (VWR) and frozen in dry-ice cooled isopentan (Sigma). 70 μm thick longitudinal section of bones were cut on cryostat (Leica), blocked with 5% goat or donkey serum and 3% BSA, and stained overnight with primary antibodies in 4^0^C, followed by 1 h staining with secondary antibodies in room temperature. Primary antibodies used for immunohistochemistry include: HO-1 (cat. SPA894, Enzo Life Sciences), CD31 (clone MEC13.3, BD Biosciences), endomucin (clone V.7C7), Sca-1 (cat. AF1226, R&D), SDF-1α (cat. sc-6193, Santa Cruz Biotechnology), CD140ß (clone APB5, eBioscience). Secondary antibodies used in the study include: donkey F(ab^-^)2 fragments (Jackson Immunoresearch) and donkey or goat whole IgG fragments (Molecular Probes) conjugated with Alexa Flours dyes or Cy3 dyes. Imaging was done on LSM780 confocal microscope (Zeiss) and analyzed in ImageJ software.

### Bone marrow transplants and chimerism analysis

Mice were irradiated with Cs-137 source with total dose 900 cGy in two equal split doses, 4 hours apart. 24 hours after the last dose mice were transplanted by tail vein injection with HSCs, or donor-derived whole BM cells in case of secondary transplantation. The donor cells were distinguished by GFP expression. HO-1^+/+^ HSC transplanted to HO-1^+/+^ or HO-1^-/-^ recipients (data in Figure 6B) were transduced with GFP coding lentiviral vectors according to previously published protocol [60]. In brief, 360 HSC were sorted to serum-free expansion (StemSpan SFEM, Stem Cell Technologies), supplemented with 20% of BIT 9500 supplement (Stem Cell Technologies), 0.1% of Ex-Cyte supplement (Millipore), 5 μg/ml Polybrene, 10 ng/ml murine stem cell factor (mSCF, Peprotech), 100 ng/ml human thrombopoietin (hTPO, Peprotech), transduced with 100 MOI lentiviral vectors. After 12 hours, all cells were transplanted together with 2×10^5^ whole BM supporter cells.

For transplantation of HO-1^+/+^ or HO-1^-/-^ HSCs (data in Figure 3N) we used mice strain HO-1^+/+^ GFP+ and HO-1^-/-^GFP+. The chimerism was checked by peripheral blood bleeding from retro-orbital sinus or submandibular vein puncture. The percentage of GFP^+^ donor cells were checked among total blood cells (CD45+), or granulocytes (CD11b+Gr-1+SSC^high^), B-cells (CD11b^-^Gr-1^-^ B220+CD3^-^) and T cells (CD11b^-^Gr-1^-^B220^-^CD3+). The positive recipients were defined as having at least 0.5% chimerism among granulocytes, what was above detection threshold based on GFP-mice.

### Transcriptome analysis by RNA-seq

To prepare libraries from small number of cells we used previously described Smart-seq2 protocol [61]. In case when 300-3000 cells were sorted (analysis presented in Figure 2 and Figure 4) RNA was isolated using the Single Cell RNA Purification Kit (Norgen Biotek). In case were 25-250 cells were available, the cells were sorted directly to lysis buffer (0.2% Triton X-100 with RNAse inhibitors), which was used for next steps of Smart-seq2 protocol (data presented in Figure 7E-J). The quality of RNA and material during preparation of libraries was checked by Bioanalyzer. The samples were sequenced on NextSeq500 (Illumina) with 75 bp single-end reads, aligned to mm10 reference mouse genome by BWA or STAR mapping software (~85% mapping efficiency with ~10-20 million uniquely mapped reads/sample). Differential gene expression (DGE) analysis was done using DEseq2 package [62] and R software environment. We analyzed differentially expressed genes only between libraries prepared in the same experimental batch. In the experiment where limited number of cells were available (data presented in Figure 7E-J), we repeated sorting the control groups and library preparation to eliminate batch effect and equal drop-out rate among libraries.

### Statistical analysis

Statistical analysis was conducted using GraphPad Prism software. Data are shown as mean ± SEM. When two groups were compared two-tailed unpaired t-test was applied. In case when more than two groups were compared, one-way or two-way Anova with Sidak or Bonferroni post-test was performed. Number of samples per group and number of independent experiments are described in the figure legends. Results were considered as significant for p<0.05 (* - p< 0.05, ** - p<0.01, *** - p<0.001).

## Supplemental Figures

**Figure S1.**
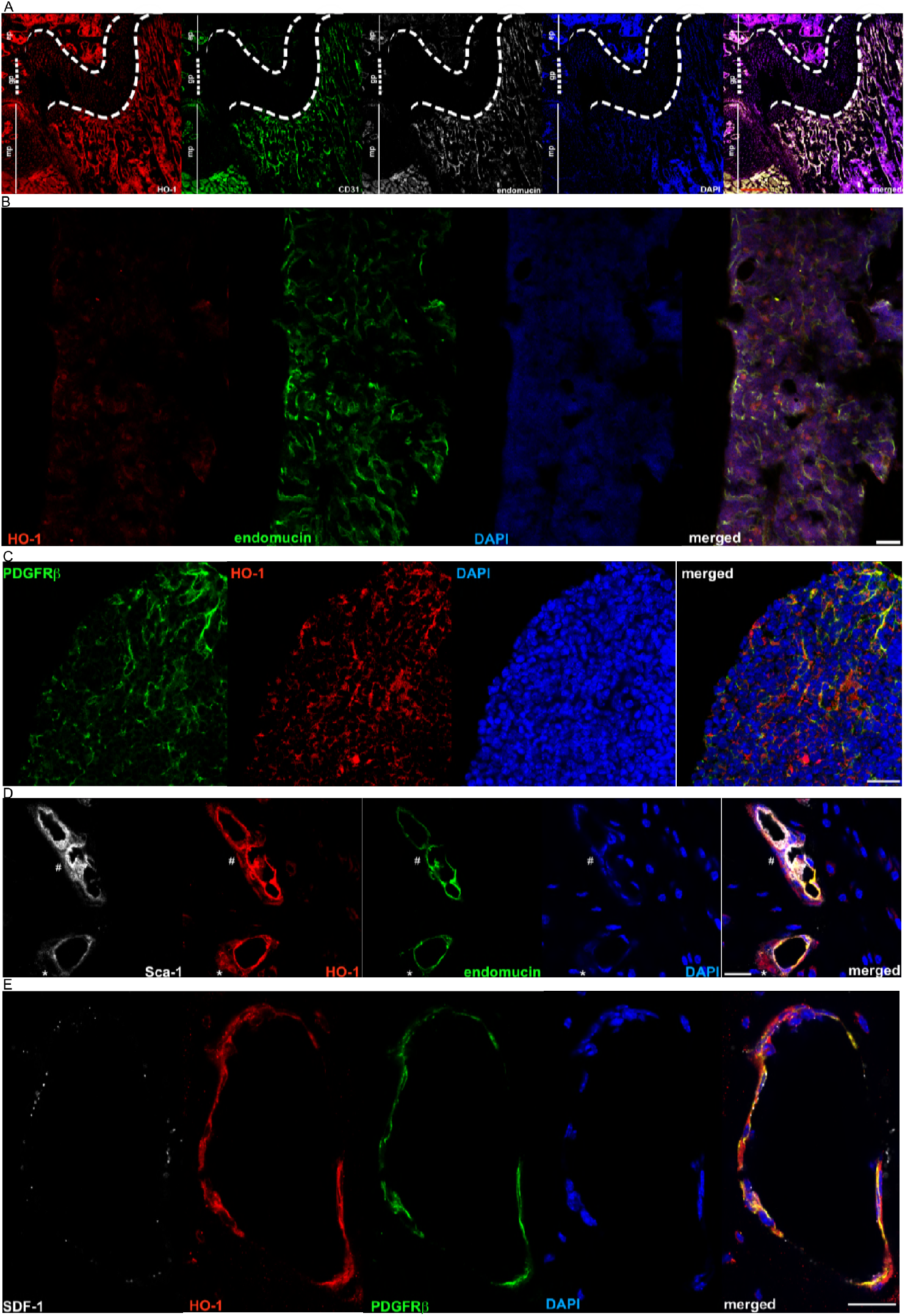
Related to Figure 1. Expression of HO-1 in BM niche. (A) HO-1 is expressed by CD31^+^ endomucin^+^ endothelial cells in metaphysis region of a tibia, bar 200 μm. (B) HO-1 is expressed in sinusoids in diaphysis region, however at lower levels, bar 100 μm. (C) PDGFRβ^+^ stromal cells in diaphysis region of the bone express HO-1, bar 20 μm. (D) Pericytes express HO-1. Part of the HO-1^+^ pericytes express Sca-1 (#), while others express no or low levels of Sca-1 (*), bar 20 μm. (E) HO-1 is expressed by PDGFRβ^+^ stromal cells. Part of HO-1+PDGFRβ^+^ cells produce SDF-1α, bar 20 μm.

**Figure S2.**
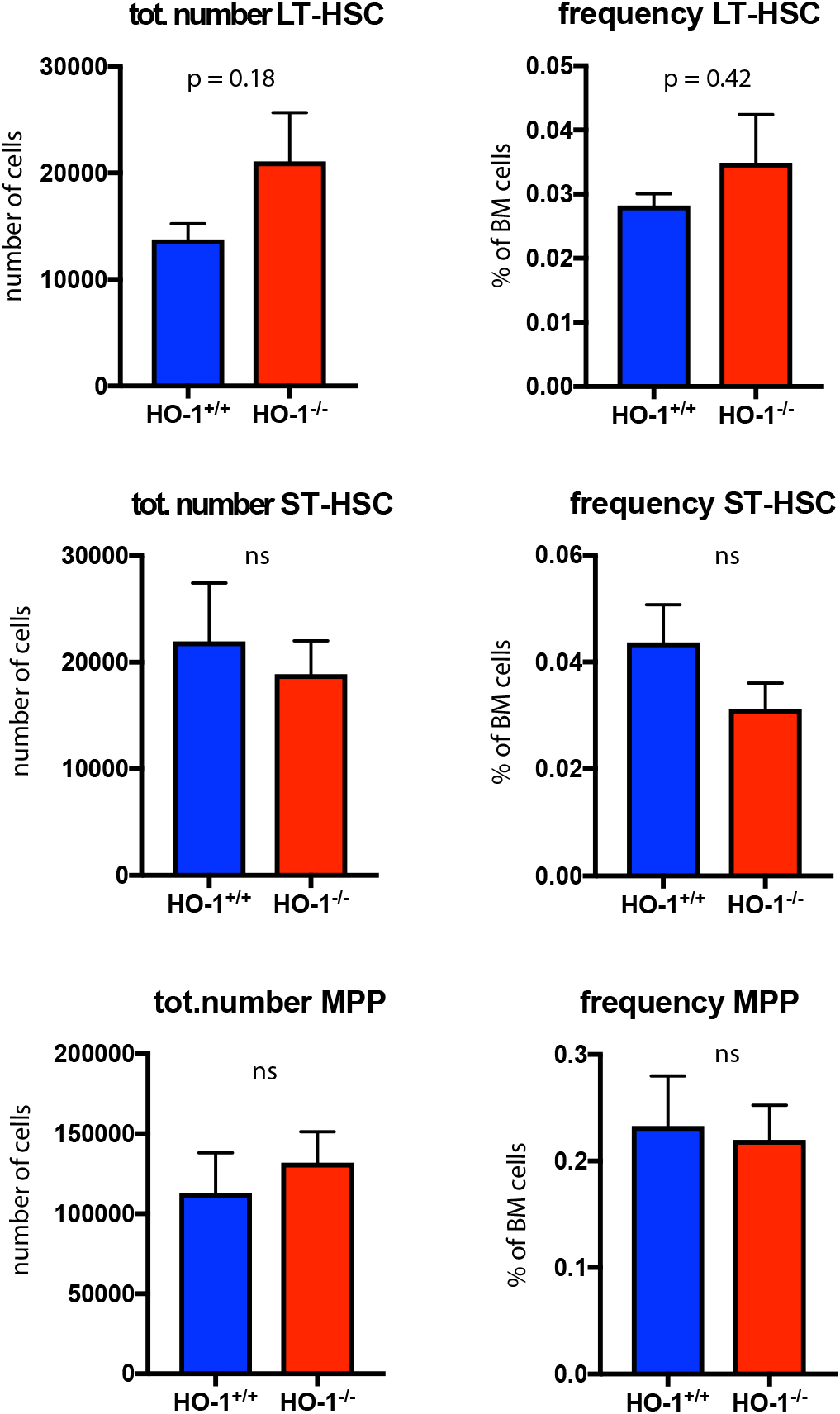
Related to Figure 3. Frequency and total number of LT-HSCs, ST-HSCs and MPPs in 12 moth old mice, n = 5 mice/ group.

**Figure S3.**
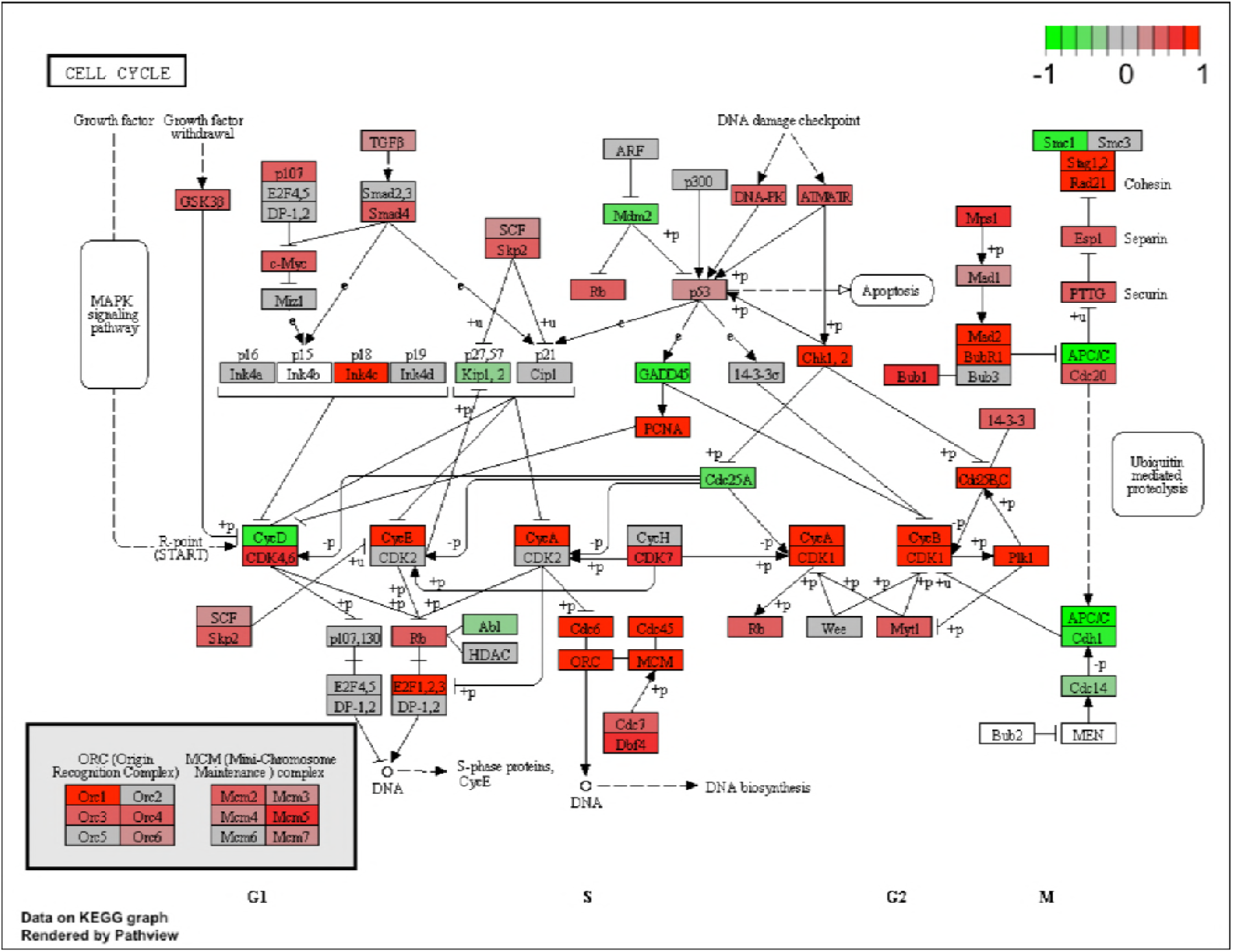
Related to Figure 4. KEEG pathway analysis showing the alterations among genes linked with cell cycle in young HO-1^-/-^ LT-HSCs.

**Figure S4.**
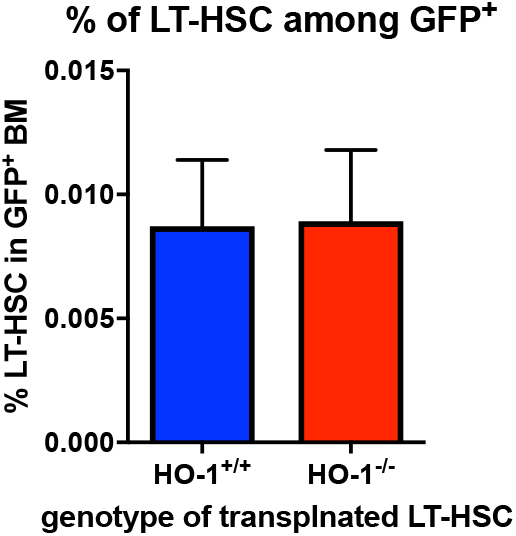
Related to Figure 7. Percentage of LT-HSC among all GFP+ donor derived BM cells in primary recipients, n = 7-8 mice/group.

## Supplemental Tables

Table S1. Related to Figure 2. RNA-seq differential gene expression analysis between HO-1^-/-^ vs HO-1^+/+^ in ECs.

Table S2. Related to Figure 2. RNA-seq differential gene expression analysis between HO-1^-/-^ vs HO-1^+/+^ in CARs.

Table S3. Related to Figure 4. RNA-seq differential gene expression analysis between HO-1^-/-^ vs HO-1^+/+^ in young LT-HSCs.

Table S4. Related to Figure 4. RNA-seq differential gene expression analysis between HO-1^-/-^ vs HO-1^+/+^ in old LT-HSCs.

Table S5. Related to Figure 4. RNA-seq differential gene expression analysis between young and old HO-1^+/+^ LT-HSCs.

Table S6. Related to Figure 4. Overlapping genes between DEGs in young HO-1^-/-^ LT-HSC and DEGs in old HO-1^+/+^ LT-HSCs.

